# Two types of hand withdraw movement to place food in the mouth mediated by somatosensation in 22-species of strepsirrhines

**DOI:** 10.1101/2022.03.13.484147

**Authors:** Louise R Peckre, Anne-Claire Fabre, Christine E Wall, Emanuelle Pouydebat, Ian Q Whishaw

## Abstract

The evolution of visual control of the hand to assist feeding by primates is uncertain but in anthropoid primates vision contributes not only to reaching for food and grasping it but also to the withdraw movement that brings food to the mouth. The strepsirrhines are a relatively large monophyletic group of Euarchontoglires near the base of the primate cladogram that are described as using vision to reach for food, but it is not known whether they use vision to assist the withdraw movement. The present study answere this question in 22 species of captive strepsirrhines from 6 of the seven strepsirrhine families, Daubentoniidae, Cheirogaleidae, Indriidae, Lemuridae, Lorisidae and Galagidae. Animals were videorecorded as they ate their normal food provisions. Dependent measures for analyses were ground withdraw movements, bringing grasped food to the mouth, and inhand withdraw movements, brining food held in the hand to the mouth, as well as the posture and head movements associated with each type of withdraw. Frame-by-frame scores from the video record showed that there were large differences between and within strepsirrhine families in these movements. Nevertheless, for all species, the withdraw movement was mediated by somatosensation, with mouth reaching and perioral contact with food determining how food was eventually eaten. Nonvisual behavior also contributed to food grasping as many species sniffed food before or during grasping. Even amongst species that made most use of the hand for their withdraws, the insectivores *Loris lydekkerianus* and *Galago senegalensis*, and herbivores, *Hapalemur simus* and *Eulemur flavifrons*, perioral contact was used to orient food for biting. The use of somatosensation and the absence of vision in mediating getting food in strepsirrhines suggests that visual mediation of the withdraw is an anthropoid innovation.

## Introduction

Primates are proposed to be special in their capacity to use vision to guide reaching and grasping of food items (Leopold et al. 2020). Both Cartmill’s (1972, 1974, 1992, 2012) visual-predation theory and Sussman’s (1991; Sussman & Raven, 1978; Sussman et al., 2013) primate-angiosperm theories posit that a stem primate evolved visual control of the hands for the capture of small food items on the terminal branches of trees. Nevertheless, these theories are unclear with respect to the details of this evolution, because nonvisual control of the hand is used with good effect for eating in many animal species (Whishaw and Karl, 2019; Iwaniuk & Whishaw, 2000; Sustaita et al., 2013) and also because the visual control of the hand is unlikely to be a unitary process. With respect to the latter point, Jeannerod’s (1981; Jeannerod et al., 1995, 1998; see also Arbib, 1981; Grant & Conway, 2019; Sartori et al, 2015) dual visuomotor channel theory proposes that visual control of the hand involves at least two neural processes. The reach directs the hand toward the extrinsic (spatial) features of a target and the grasp directs hand shaping in relation to the intrinsic (size, shape) features of a target. In addition, a food item once grasp must be brought to the mouth with a withdraw movement (Karl et al. 2018; Karl and Whishaw 2013; Whishaw and Karl 2014; Whishaw & Karl 2019). The idea that hand use for feeding is a composite 3-phase movement in turn implies that each phase has its own objective, sensory control and evolutionary history (Whishaw & Karl 2019).

Based on a study of a free-ranging population of long-tailed macaques (*Macaca fascicularis*), Hirshe et al (2022) suggest that an origin for the visual control of hand shaping for grasping may be related to the withdraw movement. They show that macaques use vision not only to appropriately grasp a food item but also to orient a food item that is grasped for subsequent appropriate presentation to the mouth. They find that vision contributions to two types of withdraw movement, the ground withdraw movement, a movement by which the hand brings a food item directly to the mouth after it is grasped, and to the inhand withdraw movement, a movement by which the hand brings a food item held in the hand to the mouth. During both withdraw movements, vision contributes to orienting food, especially food that protrudes from the hand, so that it can taken with a precise bite. The macaque withdraw movement is similar to that used by other anthropoid species including humans, suggesting that it is a behavior common to anthropoids (de Bruin et al., 2008; Karl and Whishaw,2013; Sacrey et al., 2011). The presence of visual control of the withdraw in anthropoids raises the question of whether there are anticedents in other primate species. Strepsirrhines, a relatively large monophyletic group of Euarchontoglires near the base of the primate cladogram lack the hand shaping movements featured in the visual control of precision grasps described for anthropoids but do use vision to reach for food (Bishop, 1964; Christel, 1993; Christel & Fragaszy, 2000; Macfarlane & Graziano, 2009; Marzke et al., 2015; Peckre et al., 2019; Pouydebat et al., 2008; Scott, 2019). The purpose of the present study was to examine whether vision contributes to the withdraw movement of one or more to the streptsirrhine species.

For the study, withdraw movements were scored in 22 species of strepsirrhines comprising members of 6 of the strepsirrhines families. The video recordings used for the study were those previously compiled in Peckre et al (2019a). The video recordings of feeding behavior were examined frame-by-frame and the incidence of two types of withdraw movement, ground withdraw and inhand withdraw, along with their associated head orienting and body posture were scored in relation to the putative use of vision. The rating scales for movement were designed to designate species that made little use of vision at the bottom of the scales and to describe visually mediated withdraw such as that of macaque at the top of the scales.

### Subjects

Data was collected from 84 individuals (42 female,42 male) of 22 different species of strepsirrhine primates (Table 1). The sample included six of the seven strepsirrhine families: Daubentoniidae, Cheirogaleidae, Indriidae, Lemuridae, Lorisidae and Galagidae. Greater bamboo lemurs, *Hapalemur simus*, were video recorded at Vincennes Zoo (Paris, France). Grey slender lorises, *Loris lydekkerianus*, and Senegal bushbabies, *Galago senegalensis*, were recorded at the Antwerp Zoo (Antwerp, Belgium). The other species were videorecorded at the Duke Lemur Centre (Durham, NC, USA). Animal handling was performed in compliance with the International Primatological Society (IPS) Guidelines for the Use of Nonhuman Primates in Research according to protocol #A089-14-04, approved by the Duke University Institutional Animal Care and Use Committee. For details of filming, housing conditions, and relative age, see Peckre et al (2019a).

### Food

Each animal was video recorded in its home enclosure for several days during its feeding period while eating its usual diet. The usual diet was raw pre-cut pieces of fruits and vegetables in addition to monkey chow (Labdiet Monkey Diet Jumbo Constant Nutrition and ZuPreem Primate Dry Diet). Insects were part of the diet for some species. Food items were placed on a flat surface, either on the ground level or on a raised platform, and either directly in contact with a flat surface or on a flat container (e.g., paper plate).

### Video recording

Video cameras used for diurnal species were (SONY HDR-PJ790V, full HD 1080, 24.1MP; SONY HDR-SR11, 10.2MP; SONY Handycam, HDR-PJ230, 8.9MP; and SONY HDR-CX240E, full HD 1080, 9.2MP). For nocturnal species a low-light digital video camera (SONY HDR-SR11 10.2MP) was used.

### Behavioral scoring

Four movements associated with getting food to the mouth were scored. Ground withdraw movements were scored for the posture used to pick up food and the hand movement that brought the food to the mouth. Inhand withdraw movement were scored for the posture adopted for eating and the hand movement that brought food to the mouth. Because a function of the visual contribution to the withdraw of food to the mouth in anthropoids is to place a food item in the mouth where it is taken with a single bite, this behavior was represented as the upper end of the scales for ground withdraw and inhand withdraw.

#### 1. Posture and head orientation when picking up food

The posture and orienting movement of the head and body prior to food grasping was rated on a 5-point scale:

0 – food was grasped with the mouth
1 – the nose was placed proximal to the target as the item was grasped by hand
2 – the nose was first placed near the target but withdrawn for the hand to advance
3 – the nose was first placed near the target but withdrawn at some distance
4 – the target was at arm’s length from the nose as the reach was performed

#### 2. Ground withdraw

Ground withdraw was a hand movement that brought a food item to the proximity of the mouth immediately after it was grasped. Ground withdraw of the hand was scored on a 6-point scale:

0 – hand and mouth grasped at about the same time or hand grasped or mouth reached for item as it was grasped
1 – the hand was rotated (supinated or pronated) after grasping and the mouth reached to take the item from the hand
2 – the hand made a small withdraw toward the mouth as the mouth reached to take the item from the hand
3 – the hand and the mouth toward each other was approximately equivalent
4 – the hand moved to the mouth with little movement of the head and the food item was sniffed or touched to the lips before being grasped by the mouth
5 – the hand moved to the mouth and the mouth opened to directly receive food

#### 3. Posture and orientation when taking food from the hand

Eating posture as an animal obtained a food item that was held in the hand was rated on a 5-point scale

0 – a quadrupedal posture with the back horizontal
1 – a three-point posture, with one hand on the floor and the other holding the food, with the back horizontal
2 – a two-point posture, with one or both hands holding a food item, with the back horizontal
3 – a three-point posture, with one hand on the floor and the other holding the food, with the back oblique
4 - a two-point posture with the back oblique and one or both hands holding a food item

#### 4. Inhand withdraw

Inhand withdraw movements brought a food item that was held in the hand to the mouth. Most food items held in hand were first taken from the mouth after a ground withdraw movement. Inhand withdraw movements were scored on a 6-point scale:

0 – the hand remained in place and the mouth reached to the hand
1 – the hand was rotated (supinated or pronated) as the mouth reached to the hand
2 – the hand made a small withdraw as the mouth reached to the hand
3 – the hand and the mouth moved an equal distance
4 – the hand moved to the mouth with little movement of the head and the food item was sniffed or touched to the lips before being grasped by the mouth
5 – the hand moved to the mouth and the mouth opened to directly receive food

### Statistical analyses

The results were collected using a non-probability method in which each species served as a statistical subject (Fuller, 2011). Because there were unequal numbers of animals in groups and because the number of eating movements in each of the animals differed, observations were grouped for the analyses with the result that the number of observations per species ranged from 80 to 651 (mean 322 ± 34.5 per species). Results were summarized by mean and standard error of counts of behaviors in different species and were compared using Pearson product correlations between species behaviors, with significance set at P<0.05.

## Results

### General observations

The results are based on food items that the lemurs either picked up with a hand or they picked up by mouth and transferred to one or both hands (and excluded food items picked by mouth and directly swallowed). These food items were relatively large food items, as smaller food items were more frequently picked up by mouth and then swallowed (see Peckre et al, 2019a). Frame-by-frame counts of the withdraw movements used to bring a food item were based on a total of 1,934 food items (mean 81±15.2 per species) that provided 1934 (mean 80.5±15.2) ground withdraw movements and 5530 (mean of 230.4±4.9 per species) in hand withdraw movements. Because a larger food item could be brought to the mouth repeatedly with inhand withdraw movements until consumed, there were many more inhand withdraw movements than there were food items.

Figure 1 gives the proportion of food items that were initially picked up with a hand or picked up with the mouth and transferred to the a hand. Some species displayed relatively few food item pickups using a hand (e.g., *Microcebus murinus*, *Cheirogaleus medius* and *Propithecus coquereli*), whereas other species displayed only hand food pickups (*Daubentonia madagascariensis, Hapalemur simus* and *Loris lydekkerianus*). The relative use of the mouth vs the hand was previously described by Peckre et al (2019a).

**Figure 1.**
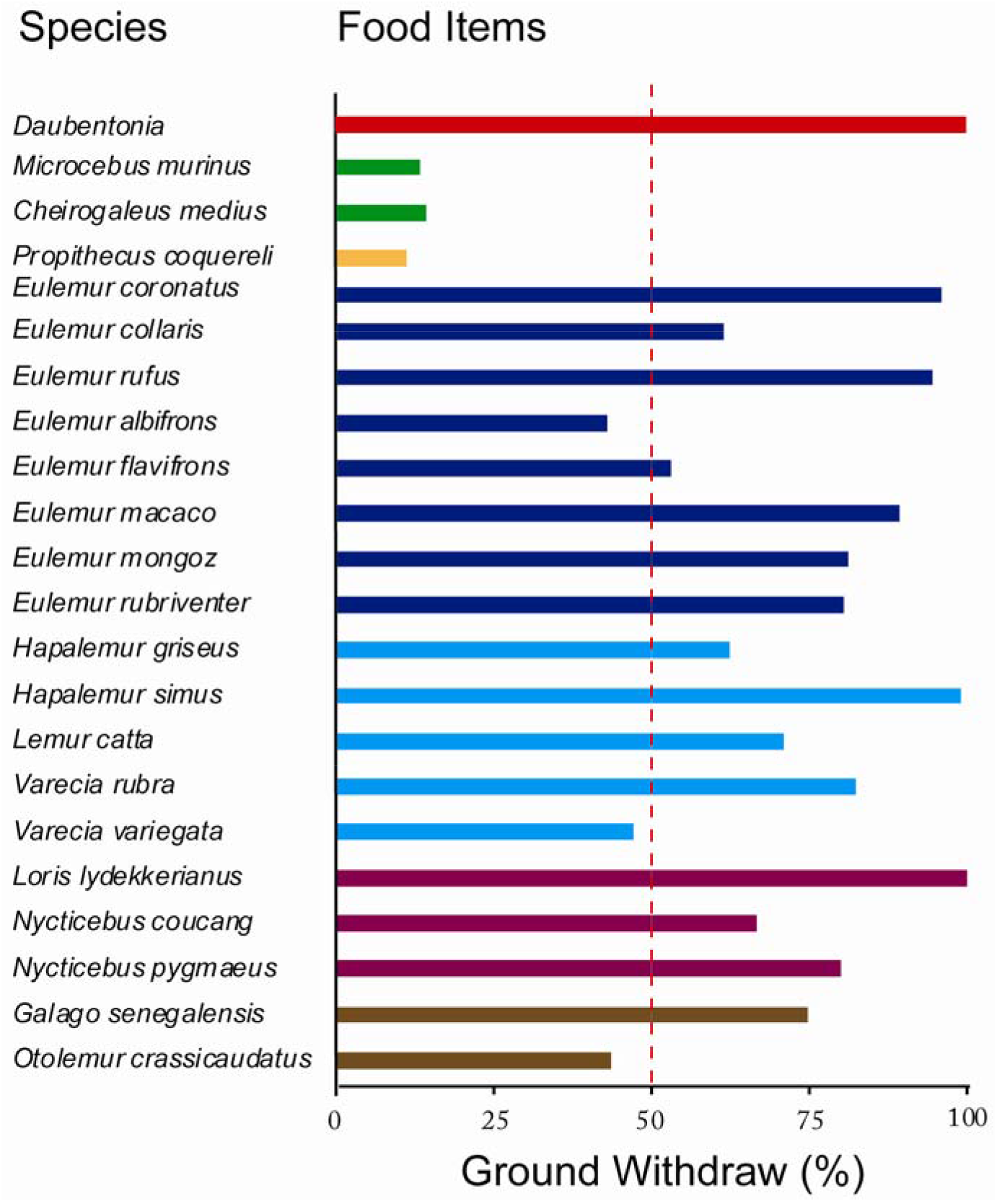
Food items picked up by hand as a percent of all food items that were handled with ground withdraw or inhand withdraw. Note: some species picked up all food items by hand (e.g., *Daubentonia*), others picked up most food items by mouth for transfer to a hand (e.g., *Microcebus murinus*), whereas still other displayed almost equal numbers of initial ground withdraw and mouth pickups (*Eulemur albifrons*).

### Relation between orienting and withdraw movements

Figure 2A describes the relationship between posture and head orientation for picking up food and associated ground withdraw ratings. Food items that were picked up by hand were always immediately brought to contact the snout/mouth by all strepsirrhine species. The relative contribution of the head and the hand to the withdraw movement depended on how proximate the head was to the food item as it was grasped. For example, if an animal was sniffing a food item as it was grasped, the hand contribution to the withdraw was small, whereas if an animal was sitting or standing back and reaching with an extended arm as the food item was grasped, hand withdraw made a large contribution to getting the food to the mouth. Many species inspected the food by sniffing, and when sniffing some concomitantly grasped the food with a hand. When the hand grasped a food item as it was being sniffed, the hand contribution to the withdraw movement was minimal because the mouth and the hand grasping the food were proximate (e.g., Daubentonia). Other species might sniff the food and then withdraw the snout so that a hand could be advanced to grasp the food, and for these species the hand made a greater contribution to getting the food to the mouth (e.g., most Lemuridae species). Other species appeared to visually identify food from a distance, or if they had sniffed the food they had withdrawn the head so that they reached from a distance. For these species, the withdraw movement was mainly a movement of the hand (e.g., *Hapalemur simus*, *Loris lydekkerianus*). Consequently, given the relationship between head orienting to identify food items and the contribution of the of the hand to the withdraw in getting a food item to the mouth, the two measures were highly correlated, r=0.95 F(1,20)=219.7, p<0.001.

**Figure 2.**
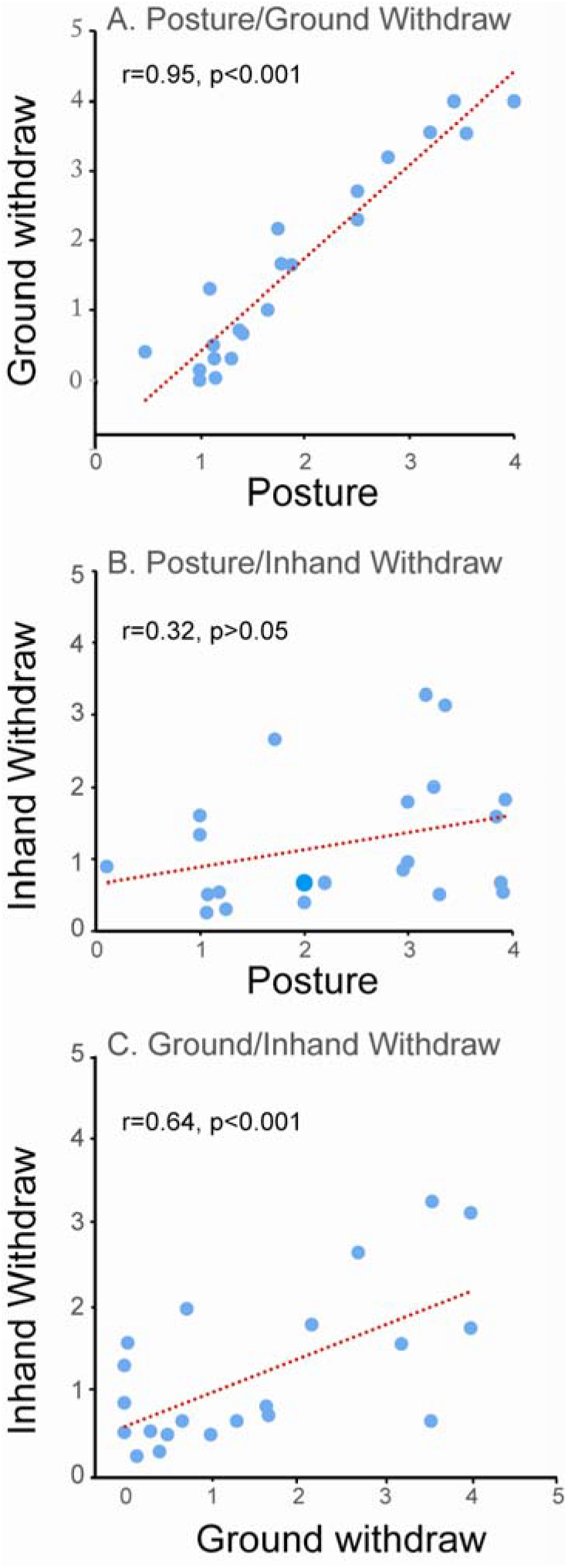
Correlations between (A) orientation and ground withdraw scores, (B) eating posture and inhand withdraw scores and (C) ground withdraw scores and inhand withdraw scores.

Figure 2B describes the relationship between eating posture and arm contributions to inhand withdraw movements. It was expected that if an animal sat in an upright position, the contribution of the hand to the withdraw movement would be greater than were the animal to crouch over the food. Nevertheless, there was no significant correlation between eating posture and the inhand withdraw movements, r=0.07, F(1,20)=0.09, p=0.76. The absence of a correlation occurred because all species made reaching movements toward the food with the mouth, both when sitting in an upright position or when hunched over the food that they were holding. In addition, some of the lemur species with long necks readily reached with the mouth for quite long distances when taking food from the hand (e.g. *Propithecus coquereli*).

Figure 2C describes the relationship between ground withdraw scores and inhand withdraw scores. The results appeared to form two clusters, some species had low scores on both measures and some species had high scores on both measures giving a significant correlation, r=0.64 F(1,20)=12.5, p=0.002. The behavior of the different species suggested that species that used orientation to sniff food before graping it or used the mouth to take food as it was grasped, also made more use of head movements to reach food in the hand.

### Species typical withdraw movements

#### Daubentoniidae

A total of 25 food item pick-ups consisting of pieces of melon, apple and banana were obtained from *Daubentonia madagascariensis*. All pick-ups were made with a hand and for all pickups, the nose was oriented proximal to the food item as the item was grasped with a hand, giving a low score of the hand contribution to ground withdraw (Figure 3A). After grasping the food, the Daubentonia always went to a nearby perch to eat. There were 55 occasions that animal raised its head away from the food held in the hand, allowing for the subsequent scoring of 55 inhand withdraw movements. On these occasions the snout moved to the food that was held in a relatively stationary hand, giving a withdraw score signifying almost no hand contribution to the withdraw. When eating food from the hand, Daubentonia adopted a variety of eating postures including being draped over a bar with the bar supporting the chest (Figure 3B), sitting on a bar with the elbows also resting on the bar (Figure 3C), or hanging by the hind feet (Figure 3D). When Daubentonia were so positioned, two kinds of eating behavior were observed. With pieces of apple, the animals mainly held the item with both hands and licked, sucked, and nibbled for long periods of time without moving their head away from the food item (Figure 3C). With pieces of banana, still enclosed in the banana peel, and with pieces of melon, they held the food item in one hand and fished for pieces of the fruit with in/out movements of the long 3rd digit of the other hand (previously reported in Lhota et al. 2009). The fishing movements consisted of many consecutive movements with a rate of 2 to 4 per sec, in which the finger-tip was inserted into the fruit and retracted into the adjacent mouth (Figure 3B). Occasionally the fishing withdraw movements would pause while the animal licked its 3rd digit, and they would then resume. Because of its idiosyncratic withdraw strategy, in which the animals seldom adopted a sitting posture and were either in a horizontal or upside-down position, eating posture score were low.

**Figure 3.**
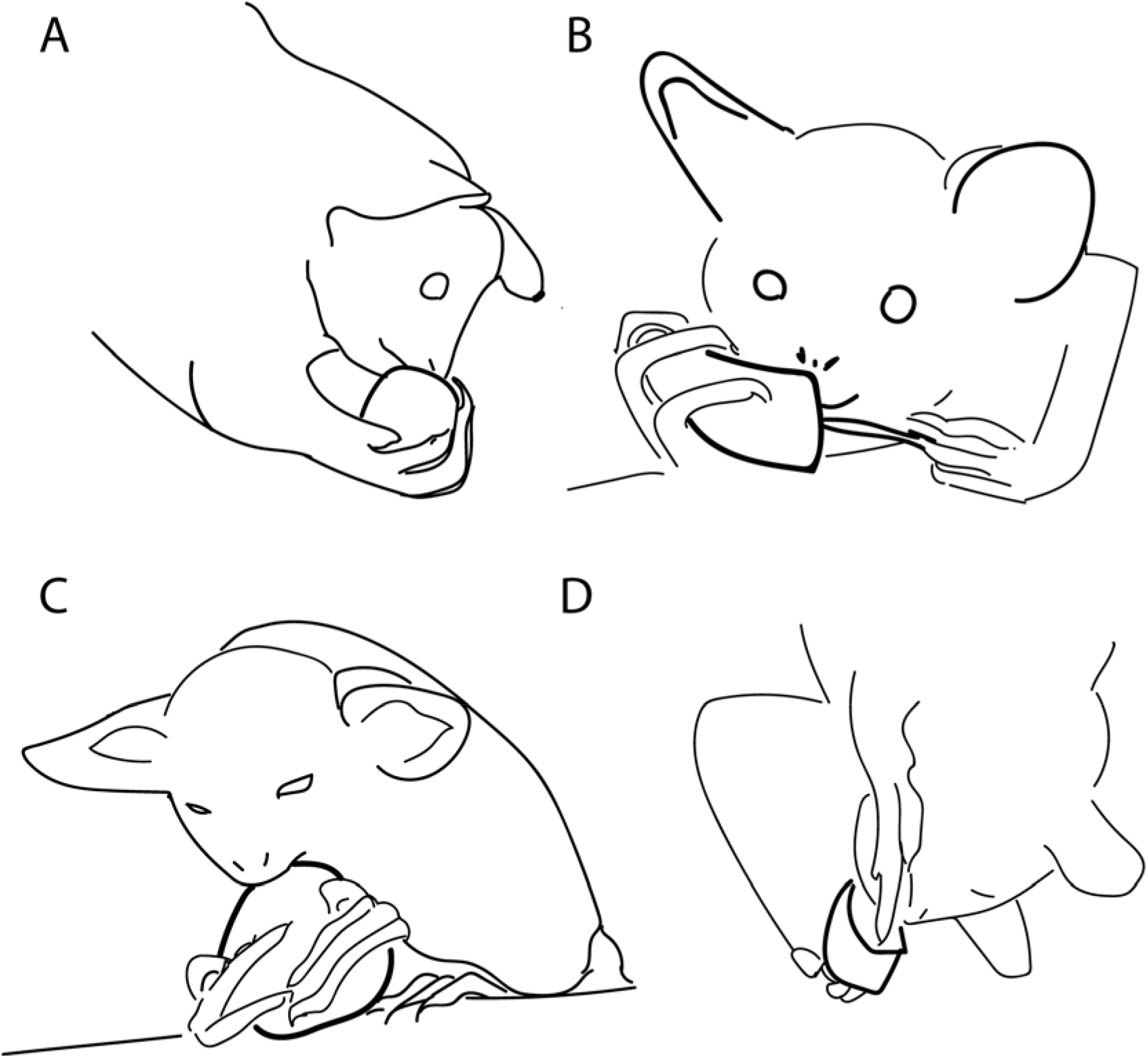
*Daubentonia madagascariensis* (A) making a ground withdraw grasp with the nose juxtaposed to the food target (apple), (B) withdrawing the contents of a banana with 3rd digit fishing movements while draped over a branch with the branch supporting the chest, (C) sitting on a pole biting at an apple that is supported by the pole and held in both hands, and (C) fishing with the 3rd digit for the contents of a melon while hanging from the hind feet.

#### Cheirogaleia

Both *Cheirogaleus medius* (observed picking up 32 food items) and *Microcebus murinus* (observed picking up 45 food items), picked up all food items using the mouth, giving orienting and withdraw scores of 0 and 0. When picking up food items with the mouth, *Microcebus murinus* adopted a quadruped stance, with forelimb placed in a wide stance and with the head extended to grasp a food item (Figure 4A). Once a food item was picked up with the mouth, *Microcebus murinus* reached and took the food from the mouth and held it in the hands for subsequent eating. In reaching for the food item in the mouth, both hands reached for the item almost simultaneously. For reaching, the hands were oriented with the palm in a perpendicular position from which they were moved to the food with an elbow-in movement. With the food held in the hands (Figure 4B), *Microcebus murinus* adopted a sitting upright oblique posture.

**Figure 4.**
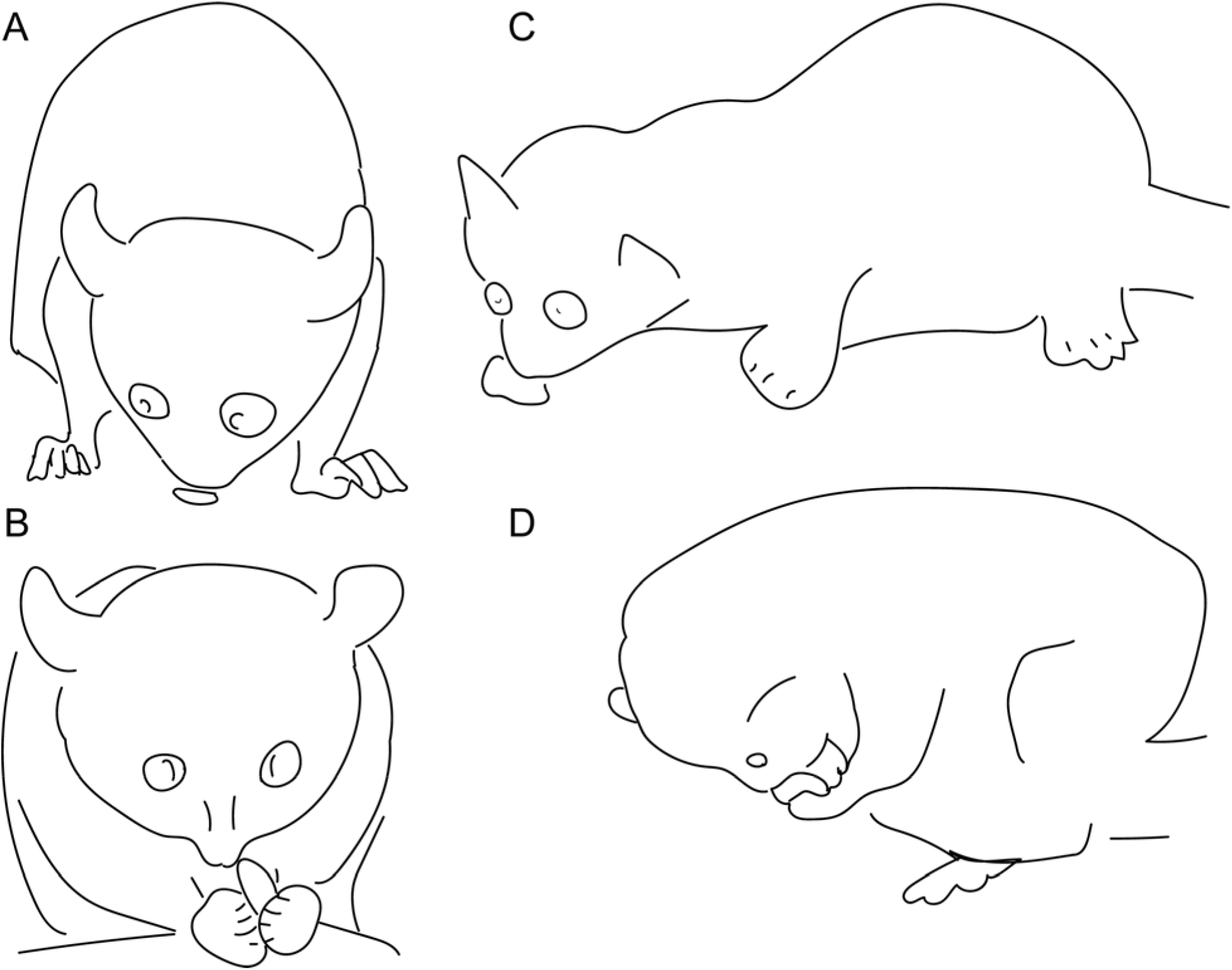
Comparison of two species of *Cheirogaleidae*. (A) *Cheirogaleus medius* stands with a wide base of support and picks up a food item by mouth, (B) that is held in both hands as the animal adopts an upright sitting posture. (C) Microcebus murinus stretches out its neck to pick up a food item by mouth, (D) that is held in both hands for eating as the animal adopts a hunched over sitting posture.

*Cheirogaleus medius* also picked up food items in the mouth but extended its head to do so (Figure 4C) and they reached and took the food from the mouth as they adopted a sitting posture. The sitting posture adopted for eating was one in which they were hunched over the food (Figure 4D), with the food item held on the substrate or with the elbows resting on the ventrum, postures that resulted in the mouth usually reaching to take food from the hand.

The bimanual food holding strategy adopted by both Cheirogaleia species resulted in most of the movement of reaching for the food being performed by movement of the mouth to the food, giving a low score for inhand withdraw movements.

#### Indriidae

*Propithecus coquereli* were observed reaching for 63 food items placed on a horizontal surface and they were also observed eating leaves (16 branches and 86 reaching events). When reaching for food items on a shelf, the animals reached for nearly all food items with their mouth (57 of 63 items). When reaching for food from the substrate, the hands usually grasped the cage wall or other objects in the cage for support. When eating leaves, the animals reached for a branch and then held the branch relatively still with one hand as they reached for the leaves to take them with the mouth. Only 7 ground withdraw movements to pick up food items using a hand were observed, all from one of the six animals. This gave an inflated ground withdraw score relative to all of the food items obtained. The reaches made by this individual were made from a distance and the withdraw mainly involved the hand coming to the mouth with the food items adjusted to be placed in the mouth after first contacting the mouth.

For inhand withdraws (n=197), the animals adopted a sitting posture with the back in an oblique orientation and with the lower arm extending at about a 90° angle from the vertically held upper arm. In-hand eating mainly involved head movements toward the food accompanied by minimal orienting of the hand, giving a low inhand withdraw score. Figure 5 illustrates the sitting posture of an animal holding a carrot (Figure 5A) and the relatively large head movement toward the carrot (Figure 5B). The carrot was not taken directly into the mouth, but was first contacted by mouth, only then to be position to the side of the mouth for biting with the molars (Figure 5C).

**Figure 5.**
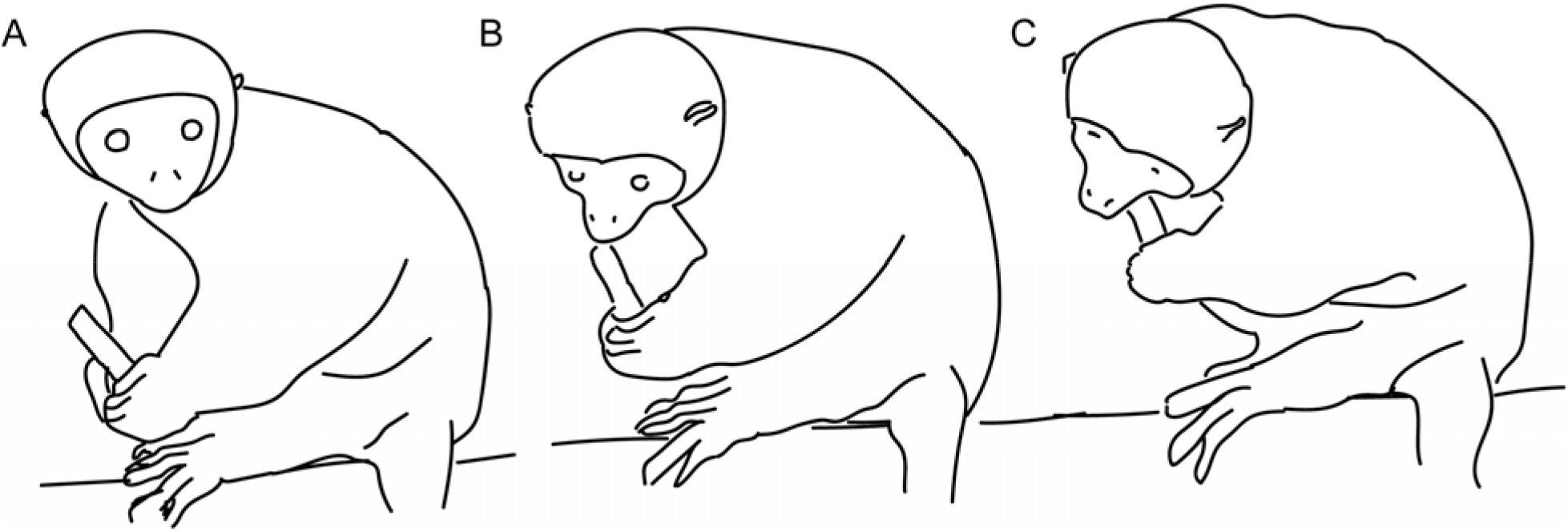
*Propithecus coquereli* (A) holds a carrot in both hands while in a sitting posture, (B) makes an inhand withdraw to bring the protruding end of the carrot to contact the tip of the mouth and (C) adjusts its head position and carrot position to bite using its molars. Note that the animal uses perioral receptors to determine how to position the carrot for biting.

#### Lemuridea

Lemuridea were divided into two groups for analyses, the Eulemurs and the other five species.

The results of the scoring of eight species of Eulemur are summarized in Figure 6. In all, 541 ground-withdraw movements (mean 60/species) and 2396 inhand withdraws movements (mean 266/species) were scored. The most frequent ground withdraw movements featured first advancing the nose toward to the target object, then grasping the object by hand, and subsequently using the mouth to take the food item from the hand. For many species the nose was proximate to the target object when it was grasped by the hand, but for some species, the distance of the nose from the target varied as indicated by scores for the orientation in Figure 6B. The cartoons in Figure 6A illustrates the orientation of *Eulemur rubriventer* that occurred with a ground-withdraw that featured the nose withdrawing but proximate to a food item so that the hand could advance to grasp, with the mouth then turning to meet the hand. Figure 6B summarizes the correlation between the orientation scores and the ground-withdraw scores for eight Eulemur species, which was significant, r(1,8)=0.94, F(1,6)=60.4, p<0.001, despite the generally low scores on both measures by all species. The correlation shows that although the nose was proximate to the food when it was grasped different species animals might raise the nose slightly more than others to allow the hand to advance onto the food.

**Figure 6.**
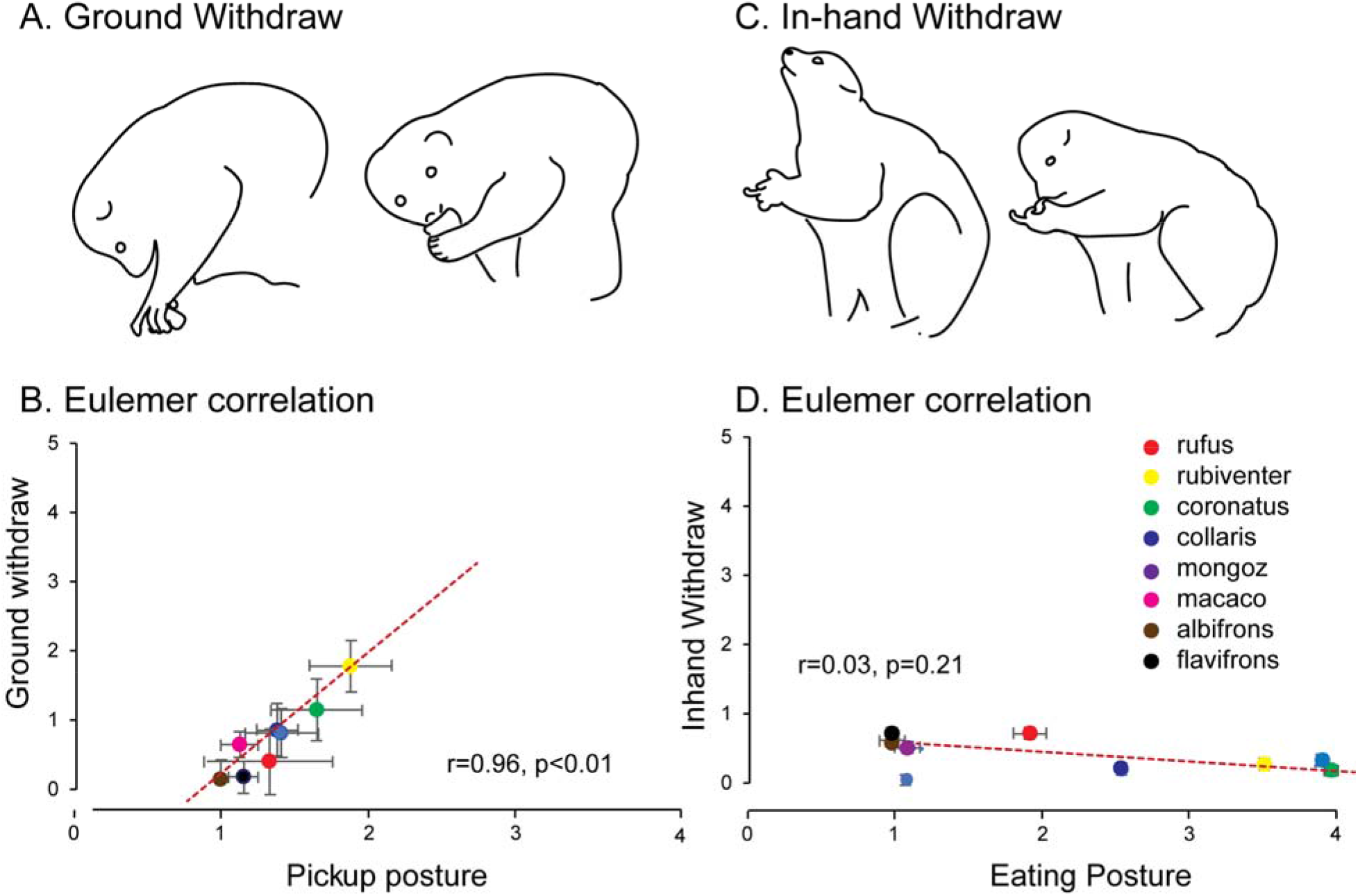
Summary of results from 9 species of *Eulemur*. (A) *Eulemur rubriventer* illustrates head orientation and ground withdraw as a food item is grasped; (B) relation between all *Eulemur* species orientation and ground withdraw scores. (C) *Eulemur rubriventer* illustrates eating posture and inhand withdraw; (D) relation between all *Eulemur* species eating posture and inhand withdraw scores.

The first cartoon in Figure 6C first illustrates a *Eulemur rubiventer* in a three-point sitting posture with an oblique back position and with its snout raised to swallow a food item. The second cartoon illustrates the *Eulemur rubiventer* lowering its snout to take a bite of food from an item held in the hand, which supinates but does not advance to meet the snout. Figure 6D summarizes the relation between all Eulemur eating posture and inhand withdraw scores. For most in-hand withdraws, the animals were in a sitting postion but the eating posture used varied by species, with some animals using a three point posture (the hand not holding the food positioned on the surface) and the other adopting a two point sitting posture, and with some animals maintaining a eating posture with the back almost horizontal and others maintaining an oblique back position. To retrieve the food from the hand all species mainly directed their mouth to the food, with the hand was held in a supinated position or else supinated on the approach of the mouth. The correlation between the scores for the eating posture of the Eulemur and the scores for their in hand withdraw movements was not significant (r=0.03, F(1, 6)=0.1, p>0.05).

The five other lemur species were scored as they picked up 411 food items and made 1,332 inhand withdraw movements (an average of 82 food items and 266 inhand withdraws per species).

Figure 7 summarizes the scores for these five lemur species each of which displayed a different patterns of ground withdraw scores and inhand withdraw. The two Hapalemur species had high scores on both withdraw movements. *Lemur catta* made more use of the hand for ground withdraw and more use of the mouth for inhand withdraw, whereas the two Varecia species had low scores for both ground withdraw and inhand withdraw movements. Figure 8 illustrates the eating posture and inhand withdraw movement for *Hapalemur simus*, a lemur that reached from a upright three-point posture resulting in an inhand withdraw that made a large contribution to getting food to the mouth. This behavior contrasts with the inhand withdraw movement for *Varecia rubra*, a lemur that frequently rested the hands holding the food on the floor and reached for the food with the mouth (Figure 8). Video 1 illustrates the use of perioral contact by *Hapalemur griseus grieus* to identify a target on a food item for biting. The animal contacts the slice of corn with its snout and then reorients the corn so that it bit off kernels from the cob (the video of this animal can be contrasted with a video of the using vision for eating a similar food item by Hirsch et al (2022)).

**Figure 7.**
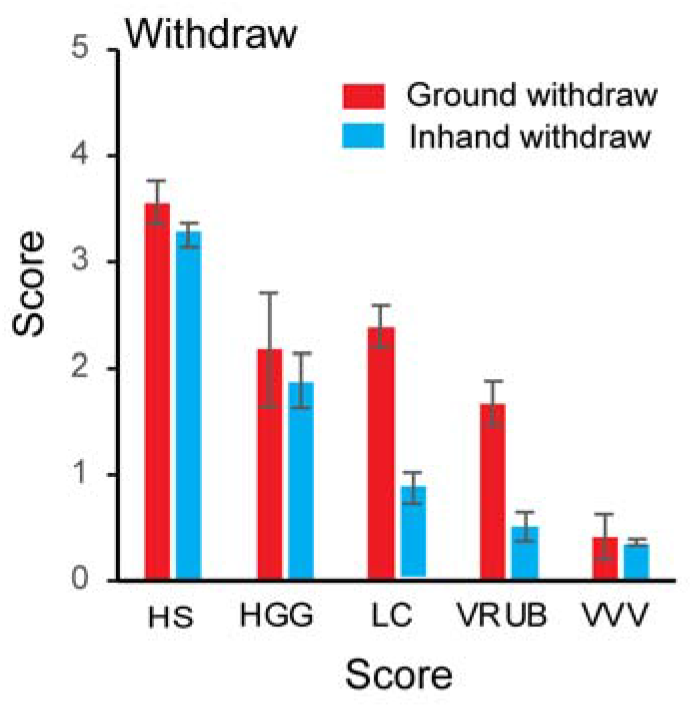
Differences in inhand withdraw by two lemur species. (A) *Hapalemur simus*, a lemur that featured an upright three-point posture and an inhand withdraw that made a large contribution to getting food to the mouth. (B) The inhand withdraw movement for Varecia rubra in which the head moves to the food and the hand makes little contribution to the withdraw.

**Figure 8.**
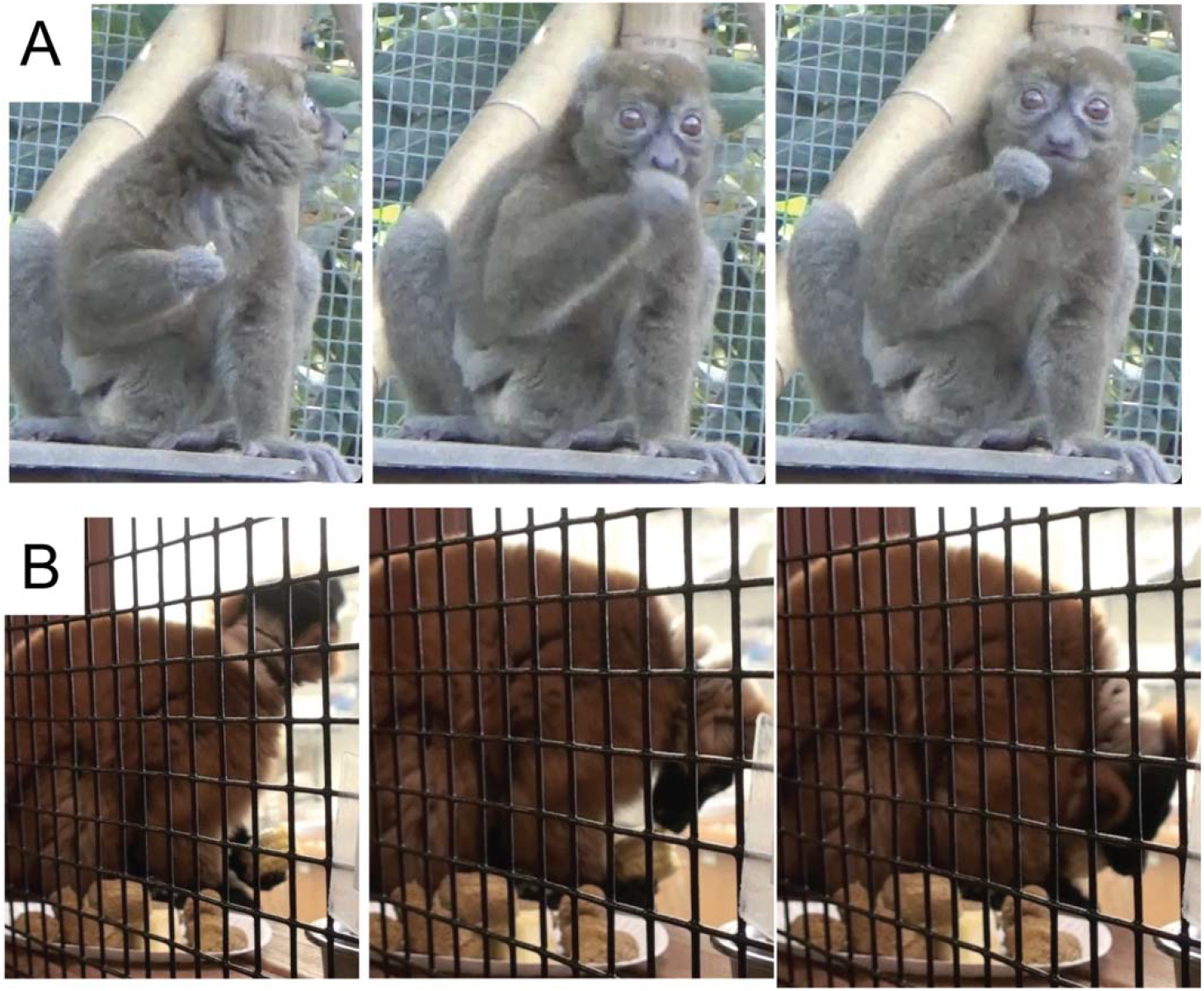
Ground withdraw and inhand withdraw scores for five lemur species.

As occurred for Eulemur species, the relationship between orientation and ground withdraw movements of the lemurs was significant, r=0.93, F(1,3)=50.5, p<0.005, but the relation between eating posture and inhand withdraw scores was not, r=0.65, F(1,3)=2.20, p=0.23. The relationship between ground withdraw scores and inhand withdraw scores was also significant, r=0.89,F(1,3)=12.5, p=0.038.

#### Lorisidea

A summary of behavioral rating scores for *Loris lydekkerianus*, *Nycticebus coucang* and *Nycticebus pygmaeus* is shown in Figure 9. Measures were obtained from 209 food items pickups that provided 417 inhand ground withdraw movements from three loris species (averages of 70 and 139 per species). The behavior of *Loris lydekkerianus* was different from the other two species. *Loris lydekkerianus* nearly always used a single hand to pick up a food item and did so at a distance after making a few orienting (lateral and back and forth) head movements to fixate the target. Its ground withdraw movement brought the food item to the snout with little concomitant head movement. After an item contacted the snout, it was either discarded or the animal sat back with the item still in its hand to eat. When in a sitting posture, the loris did make head turning movements to take the food item from its hand into its mouth and so it receives lower inhand withdraw score than ground withdraw score. *Nycticebus coucang* and *Nycticebus pygmaeus* were more likely to pick up food items with the mouth and then transfer the items to one or both hands for holding for eating. Both species used head movements to retrieve the food from the hand, but their eating posture was different. Nycticebus pygmaeus would sit to eat in a three point or two-point posture with the back in a oblique orientation. Nycticebus coucang held its elbows against the torso and arched its head underneath its torso to retrieve food from the hand with the mouth.

**Figure 9.**
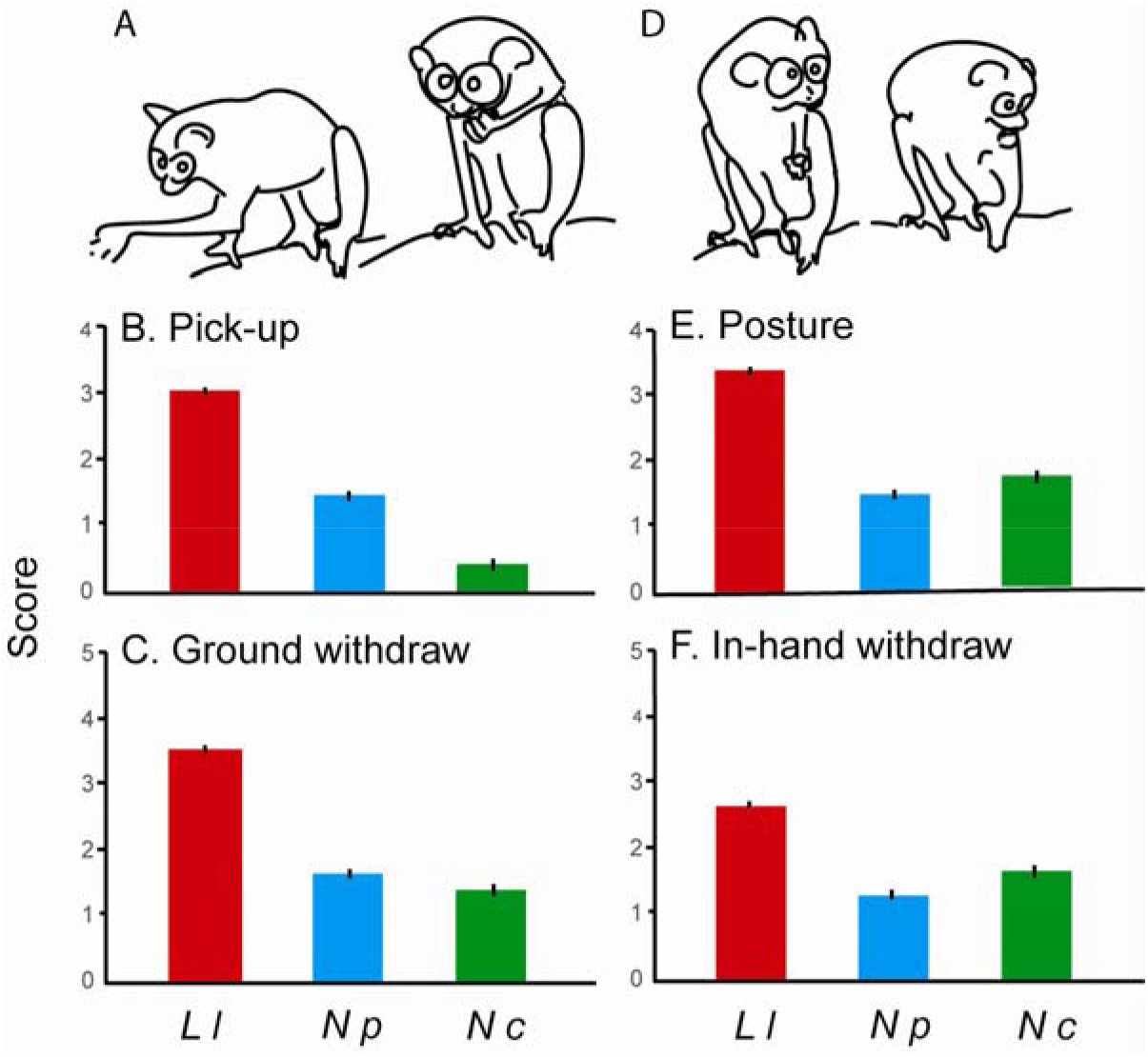
Ground withdraw and inhand withdraw for three species of Lorisidea. (A) *Loris lydekkerianus* makes a ground withdraw in which most of the movement is made by the hand. (B) Orient scores and (C) ground withdraw scores for three Lorisidea species. (D) *Loris lydekkerianus* features a 3-point oblique sitting posture from which both the head and hand contribute to the withdraw. (E) Eating posture and (F) inhand withdraw scores for three Lorisidea species.

#### Galagidea

Data were obtained from two species of Galagidea, *Otolemur crassicaudatus* (18 food items and 74 inhand withdraw observations) *andcGalago senegalensis* (302 food items and 279 inhand withdraw observations). Otolemur made all food pickups by mouth (mean score of 0) whereas Galago made about half of its pickups by mouth and the others by hand. Even when picking up food by hand, Galago made orienting movements of the nose so that the target food item was almost touched, but it often then sat back and reached for the food by hand. Figure 10 summarizes in-hand withdraw movements and eating posture of the two Galago species.

**Figure 10.**
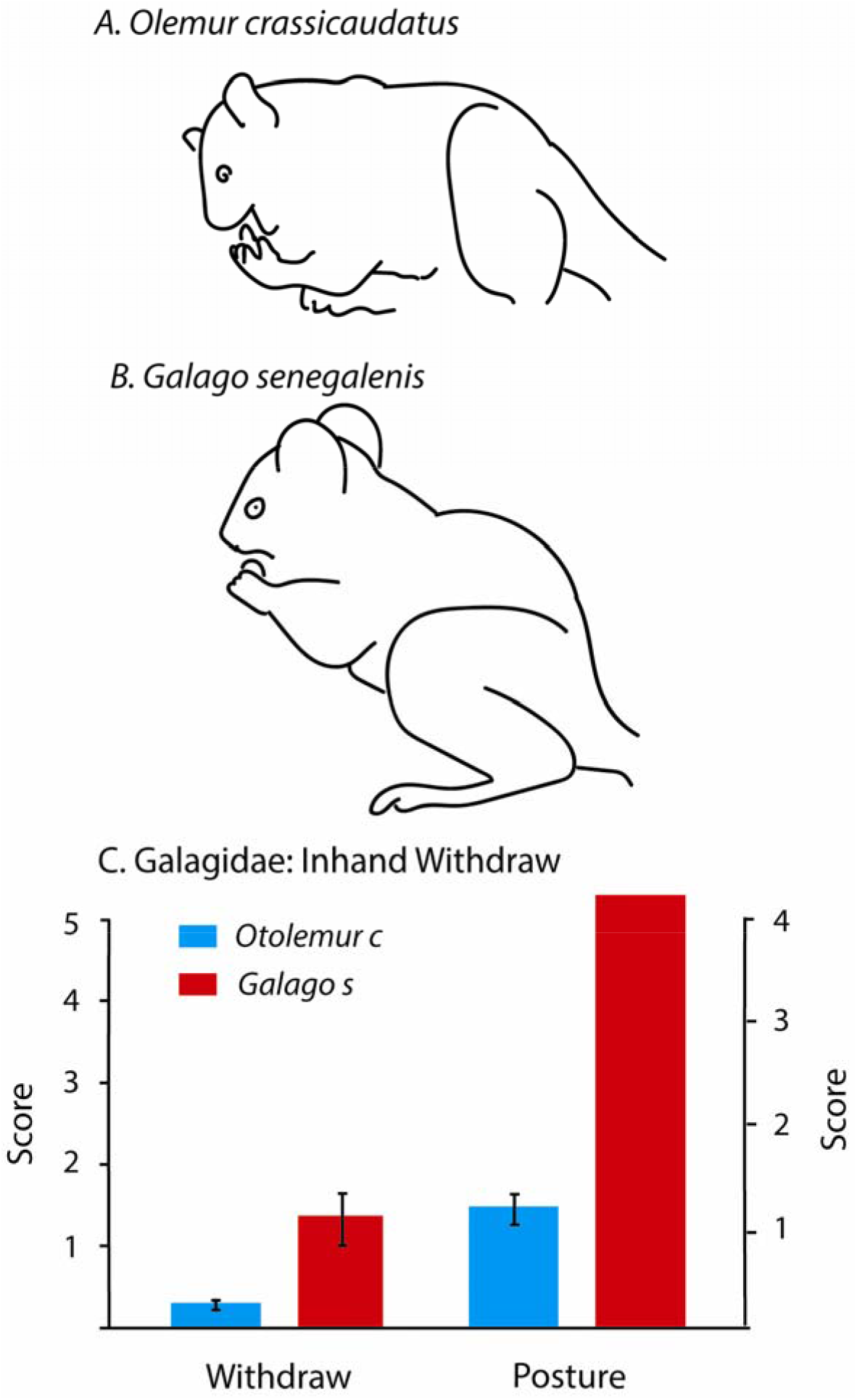
The eating posture of two species of Galagidae. (A). *Otolemur crassicaudatus* sitting posture was on the haunches with the torso horizontal. (B). *Galago senegalensis* sitting posture was on the haunches holding food items in one or both hands with the torso oblique. (C) For both species inhand withdraw consisted mainly of the mouth orienting to the food item giving low scores for movement of hands during the withdraw.

The eating posture of *Otolemur crassicaudatus* involved sitting on the haunches with the torso horizontal (Figure 10A). The eating posture of *Galago senegalensis* consisted mainly sitting on the haunches holding food items in one or both hands with the torso oblique (Figure 10B). For both species inhand withdraw consisted mainly of the mouth orienting to the food item giving low scores for inhand withdraw (Figure 10C).

### Comparison of species with high ground withdraw scores

There were differences in the ground withdraw movements, even amongst species with the highest ground withdraw scores. A representative ground withdraw by *Loris lydekkerianus* (mean withdraw score 4) is shown in Video 2. The loris reaches out to almost touch the target food item with its nose, but after this movement is completed and the head withdrawn, it then reaches for and grasps a selected food item with its hand. The hand brings the food item directly to the mouth, but it does not take the food item into the mouth. Only after it sits back it sniffs the food does it take a bite. A representative ground withdraw with a captured worm is shown in Video 3 for *Galago senegalensis* (mean withdraw score *3.54*). The reach is made from a distance and the worm is brought directly back to the mouth but only taken into the mouth after a contact. A representative ground withdraw followed by an inhand withdraw is shown for *Hapalemur simus* (mean withdraw score 3.2) in Video 4. The food item is visualized as the reach begins but vision is disengaged before the grasp is completed. The withdraw brings the food item to the front of the mouth and then after contact, the food item is moved to the side of the mouth for biting with the molars. For a subsequent inhand withdraw, the hand and the head make almost equal contributions and the food taken into the mouth following perioral contact.

## Discussion

The objective of this study was to investigate the contribution of vision to the withdraw movement, by which food is transferring from the hand to the mouth, in strepsirrhine primates. The assumption underlying the analyses is that if vision contributed to the withdraw, that would be signified by the behavior of looking at an item once grasped in the hand to ensure its subsequent accurate placement in the mouth. The withdraw was investigated during normal feeding in captive animals comprising 22 species from six families. Rating scales evaluated orienting and ground withdraw movements and orienting and inhand withdraw movements, which transferred food held in a hand to the mouth. There were large differences in the incidence and the form of withdraw behavior within and between strepsirrhine families. Nevertheless, there was no evidence that vision was involved in the guidance of the withdraw movement of any species. No species directly placed food into the mouth and all used perioral/perinasal contact to determine the edibility of a food item and to assist in orienting food items for biting. Both kinds of withdraw movement were minimized by advancing the mouth to grasp the food rather than bringing the food by hand to the mouth. Species that made most use of the hands to withdraw food included insectivores and herbivores, suggesting that somatosensory withdraw strategies are usefully applied to a wide variety of foods.

The evidence that anthropoid primates use vision to guide the withdraw movement of bringing food to the mouth is that they look at a food item as it is grasped, adjust the orientation of a food item in the hand before or during the withdraw, and after then looking away, directly place the item into the mouth where it is taken with a single bite (Hirsche et al, 2022). There was no evidence that any strepsirrhine species used these strategies to orient food to the mouth. Strategies that the strepsirrhines did use in getting food into the mouth included bringing the mouth to the food and then investigating the food by mouth touching, sniffing, or licking it before biting, or taking it with the mouth and then adjusting its position with a hand for biting. The ratings of head orientation to a food item that was to be picked up with a ground withdraw movement, indicated that for most species for most reaches, the nose was brought proximal to the food item suggesting that olfaction was contributing to item identification. This snout/hand proximity resulted in the item being transferred from the hand to the mouth almost immediately, precluding the use of vision. When food items were held inhand, most species oriented the head to the food held in the hand, again minimizing the role of hand movements for the withdraw. For both ground withdraw and inhand withdraw movements, food items were usually investigated perorally before transfer to the mouth. These observations suggest that all animals used nonvisual cues to guide food items as do may other animal species. For example, tree kangaroos (*Dendrolagus*) use tactile cues on the mouth in association with head movements to guide vegetation into the mouth from the hand (Iwaniuk et al., 1998), rodents use vibrissae cues and head movements to guide food to the mouth (Whishaw & Coles, 1996; Whishaw et al., 1998, 2018, 2020) and the gray short-tailed opossum (*Monadelphis domestica*) uses a wide-open mouth to receive a prey item from the hand (Ivanco et al., 1996),

Most of the strepsirrhine species adopted a sitting posture when handling food and eating it and most held the food in one hand when eating. A sitting posture for eating appears to be a featured posture in euarchontoglires, including rodents, strepsirrhines, and anthropoids (Reghem et al, 2011; Hirsche et al, 2022; Whishaw et al, 1988). The use of a sitting posture along with one hand eating could contribute to food visualization, but there was no evidence that animals adjusted the orientation of food that they held in their hand. Amongst the species that did sit upright and did hold food in one hand, all always explored the food using perioral receptors before biting, suggesting that they used touch for food orientation. A few species preferentially used two hands to hold food (e.g., *Microcebus murinus*), hunched over their food (e.g., *Cheirogaleus medius*), or had an ideosyncratic eating strategy in which the mouth and food were always in close contact, (e.g., *Daubentonia*), all strategies that limit the contribution of vision to the withdraw. Humans and macaques are also reported to blink as part of their disengage after visualizing a food item during the withdraw (de Bruin et al., 2008; Hirsche et al, 2022; Sacrey et al., 2011) but there was no evidence that any of the strepsirrhines blinked in conjunction with any portion of their withdraw.

Previous study of a withdraw movement in macaques and humans have compared how small and large food items are transferred from a hand to the mouth. For small items held in the hand, touch perception of the object by the fingers likely provides all of the information necessary to accurately postion an object in the mouth. Experimental studies with visually occluded humans show that the fingers can be accurately directed to targets on the body (Edwards et al., 2005) and the hand can accurately place or take food from the mouth (de Bruin et al., 2008; Karl et al., 2012; Sacrey et al., 2011). If a food item is large and protrudes from the hand, however, and if the distal portion of the food is to be placed in the mouth, somatosensory information from the hand is insufficient to direct the distal end of the food item to a mouth target. Macaques and humans deal with protruding food items by visualizing the distal portion of the food to calculate the trajectory of the item into the mouth (Hirsche et al, 2022; Whishaw et al, unpublished). An analysis of how the strepsirrhines dealt with food items of different sizes could not be similarly investigated due to their propensity to pick up smaller items in the mouth, presumably because their whole hand grasping strategies were not adequate for grasping small food items (Peckre, 2019a), and because all items that they did bring to the mouth with a hand were first touched to the mouth before being oriented for biting.

A surprising aspect of the behavior of the strepsirrhines was their olfactory investigation of food before they grasped (see Nevo and Heymann, 2005). Even species that often reached for food from an arms length, including *Loris lydekkerianus* and *Galago senegalensis*, often also advanced their nose in proximity to the food prior to reaching, suggesting that they were making some use of olfaction to identify food items. In this respect it is interesting that *Galago senegalensi* reached from a distance for worms that were moving but usually sniffed other food items before reaching, suggesting that sniffing was not obligatory. Olfactory investigation may have occurred because a variety of food stuffs were concurrently present at a feeding site and the animals were searching for more favored food items. Nevertheless, the food that they were given was usual fare and so it might be expected that they would be able to visually identify preferred items. Given that many of the strepsirrhine species were nocturnal and generally have a single cone photoreceptor for short wavelengths of light, lack a dense phorecoeptor region for detail vision, and have 6 to 10 times less visual acuity than anthropoids, deficient visual acuity may promote use of olfaction to assist with food discrimination (Kirck, 2004; Veilleux and Kirk, 2009).

A number of strepsirrhine species were notable for the way in which they used a hand for withdraw movement. *Hapalemur simus* frequently made reaches at arms distance from a food item and then withdrew the food item directly to the mouth. They nevertheless visually disengaged an object at the point of grasping and only when the food item was touched to the mouth was the item oriented for biting. Furthermore, their inhand withdraw movements were similarly made mainly with the hand, but in doing so they did not look at the hand during the withdraw and again they used perioral contact to assist in transferring the item into the mouth. *Galago senegalensis* distinguished nonlive and live food items by sniffing nonlive food items and often taking them by mouth while reaching for live worms that they withdrew directly to the mouth. Again, once the hand reached the mouth a food item was not taken directly into the mouth but was first positioned using perioral contact. Indeed, it would be difficult for them to make a visual calculation given that a worms protruded from the hand was usually still moving. When making inhand withdraw movements, *Galago senegalensis* did use head movements to reach for the food and prioral contacts to facilate food transfer to the mouth. *Loris lydekkerianus* consistently made more extensive use of the hand for ground withdraw movements than did other strepsirrhines (see also Peckre, Fabre et al. 2019). However, as did the other strepsirrhines they did assist inhand withdraw movements by additionally reaching with the mouth. Although the versatile use of the hand by *Loris lydekkerianus* has been noted previously (for a review, Nekaris, 2005) there was no indication that they used vision to assist in their withdraw movements. *Loris lydekkerianus* was notable in holding a food item adjacent to the snout for a considerable time after a withdraw, which along with the hand positioning movements they made near the mouth suggested that they were using perioral cues to assist in food positioning.

A caveat relevant to the methods used in the present study is that a definitive description of vision use in food handling would require the use of eye tracking glasses. For humans, the use of eye tracking glasses show that the timing and duration of food visualization is related to orienting food for withdrawal to the mouth (de Bruin et al., 2008; Sacrey et al., 2011). The same studies report that after visualizing food for orientation, the subjects blink as they disengage as do macaques (Hirsche et al, 2022). Future work could use eye tracking glasses to confirm that the withdraw strategies used by strepsirrhines were nonvisual. The methodology of the present study was also opportunistic as the animals were eating their usual fare. Future work could designate small and large food items in a more definitive experimental design. For example, *Loris lydekkerianus* has been reported to pick up some items as small as ants (Nekars, 2005), but no such food items were used here. Finally, some aspects of the withdraw movement, such as the reorienting of food as it contacted the mouth, was difficult to describe from the few video frames in which the behavior occurred, and so future work could use higher frame rate video recording.

In conclusion, an extensive examination of thousands of withdraw movements made by strepsirrhines provided no evidence that any species used vision to assist in directing food items into the mouth. The observations did reveal that strepsirrhines are extremely versatile in using their hands to get food to the mouth despite their varied feeding proclivities as insectivores or herbivores. It is interesting that amongst the many visual system difference in strepsirrhines and anthropoid primates, strepsirrhines have a smaller agranular frontal cortex area than do anthropoids and may not have homologues to anthropoid areas 9, 12/47, 46 and 10, regions involved in object and spatial working memory of they type that might be required for visually-based food size and orientation calculations (Goldman-Rakic, 1992; Preuss and Goldman-Rakic, 1991). The present findings are consistent with the idea that the evolution of the visual control of feeding was not a singular event but evolved in stages along with many other enabling behavioral and physical modifications associated with the visual guidance of hand movements.

## Video captions

Video 1. *Hapalemur griseus grieus* inhand withdraw. The animal uses perioral contact to identify and orient a slice of corn so that it can take kernels from the cob.

Video 2. *Loris lydekkerianus* ground withdraw. First, the loris almost touch the target food item with its nose. Second, the head is withdrawn. Third, it reaches for and grasps a selected food item with its hand. Fourth, the hand brings the food item directly to the mouth, but it does not take the food item into the mouth. Fifth, only after it sits back it sniffs the food does it take a bite.

Video 3. *Galago senegalensis* ground withdra). The reach for a worm is made from a distance and the worm is brought directly back to the mouth but only taken into the mouth after a contact.

Video 4. *Hapalemur simus* ground withdraw and inhand withdraw. For the ground withdraw the food item is visualized as the reach begins but vision is disengaged before the grasp is completed. The withdraw brings the food item to the front of the mouth and then after contact, the food item is moved to the side of the mouth for biting with the molars. For the inhand withdraw, the hand and the head make almost equal contributions and the food taken into the mouth following perioral contact.

